# Cognitive-Perceptual Style on the Autism-Schizotypy Continuum Shapes Surprisal Sensitivity

**DOI:** 10.1101/2025.08.19.670092

**Authors:** Merle M. Schuckart, Sarah Tune, Sandra Martin, Gesa Hartwigsen, Jonas Obleser

**Affiliations:** Department of Psychology, University of Luebeck, Germany; Center of Brain, Behavior and Metabolism, University of Luebeck, Germany; Research Group Cognition and Plasticity, Max Planck Institute for Human Cognitive and Brain Sciences, Germany; Wilhelm Wundt Institute for Psychology, Leipzig University, Germany

**Author notes:** Merle M. Schuckart, Jonas Obleser. **Email:**.

**Keywords:** language, autism, schizotypy, predictive processing, ageing

## Abstract

How the brain weighs prior knowledge against incoming evidence varies systematically across individuals – and this variation may lie at the heart of an Autism-Schizotypy Continuum (ASC) in the healthy population. In a large lifespan sample (N = 340; age 18–82), we used the Schizotypal Personality Questionnaire and a refined Autism Spectrum Quotient model to position individuals along this continuum, then asked how their placement predicts sensitivity to lexical surprisal during self-paced reading. First, perceptual style is not fixed: older adults showed reliably less schizotypal profiles, making the ASC a developmentally dynamic dimension. Second, ASC position interacted with age to modulate surprisal sensitivity across 160,000+ word-level reading times: autism-like profiles yielded stronger disruption by unexpected words, an effect that grew across the lifespan. We interpret this as reflecting a tightening of predictions formed from linguistic context – the less schizotypy-like the profile, and the older the reader, the more strongly incoming evidence is weighted against expectation. These effects remained robust under cross-validation. Our findings establish the Autism-Schizotypy Continuum as a dynamic, lifespan-sensitive framework for understanding how individuals differ in their responsiveness to incoming linguistic evidence.

**Significance Statement:** Individual differences in perception are often framed as a trade-off between prior knowledge and sensory input, yet how these differences evolve with age and shape real-time language processing remains unclear. In a large adult lifespan sample, we show that cognitive-perceptual style shifts systematically with age and predicts sensitivity to lexical surprisal during reading. Our findings recast the Autism-Schizotypy Continuum as indexing sensitivity to incoming evidence and highlight the need for lifespan-sensitive, individualized models of predictive processing.

## Introduction

Our perception is shaped both by the sensory stimuli we encounter and the cognitive state we are in. From a Bayesian perspective, it arises from a dynamic balance between prior expectations and incoming sensory evidence, and differences in their relative weighting can manifest in distinct perceptual outcomes (see Fig. 1a).

**Figure 1.**
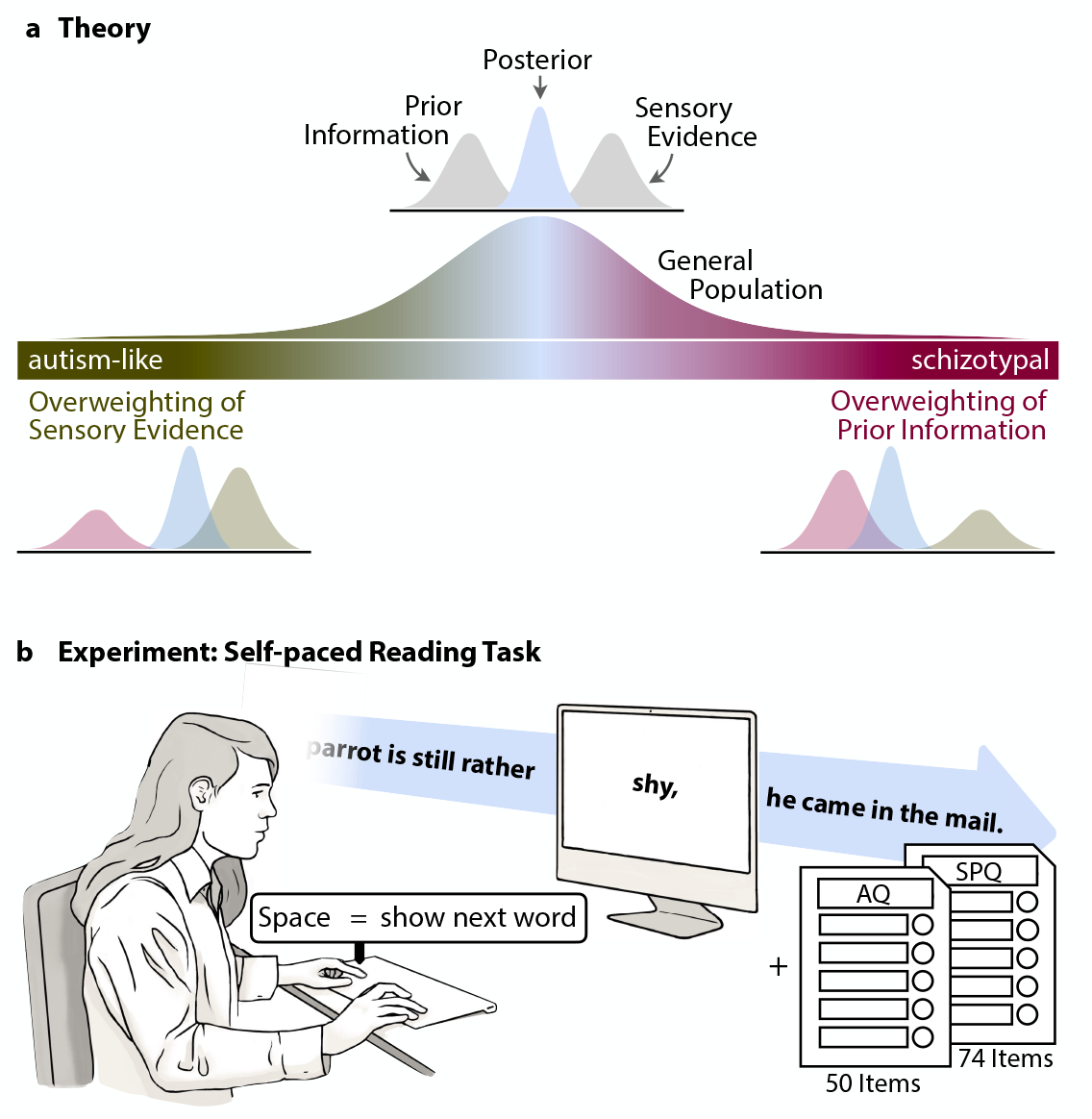
Theoretical background of the Autism-Schizotypy Continuum and design of the present experiment. a) Cognitive-perceptual style along the Autism–Schizotypy Continuum (ASC). b) Participants (N = 247) performed 2–3 blocks of a self-paced reading task. Cognitive-perceptual style was assessed using the AQ and SPQ to derive individual ASC scores.

This framework raises three central questions: (i) First, how can we capture inter-individual differences in the weighting of prior knowledge versus sensory evidence? (ii) How do such differences vary across the adult lifespan, given age-related changes in sensory precision and cognitive flexibility (1–5)? (iii) Can these differences be leveraged to explain variability in how individuals deal with the uncertainty inherent in the language we encounter?

One candidate framework for capturing such inter-individual differences is the *Autism-Schizotypy Continuum* (ASC; also referred to as the ASD-SSD continuum) which characterises variation in cognitive-perceptual style in neurotypical individuals (6–9). Specifically, the ASC has been proposed to reflect individual differences in the balance between prior knowledge and sensory evidence during perceptual inference (6–8). Within this framework, individuals vary along a spectrum from more sensory-driven (autism-like) to more prior-driven (schizotypal) inference (see Fig. 1a). The Autism Spectrum Disorder (ASD) end is associated with relative overweighting of sensory evidence and heightened attention to detail, supposedly linked to altered prediction error signalling or reduced top-down feedback. In contrast, the schizotypal end is associated with increased susceptibility to hallucinations and delusions, heightened cognitive disorganisation (10– 12), and a strong reliance on prior knowledge (13), all consistent with generally enhanced top-down influences on perception. Accordingly, an individual’s position on the ASC should show behavioural and neural correlates in perceptual inference tasks (6, 8).

To date, most studies employing the ASC have relied on relatively homogeneous samples of young adults (except 6, 7). This has not only limited generalisability but also has hitherto precluded an understanding of potential lifespan dynamics captured by the ASC construct.

Here, we address this gap by combining a large adult lifespan sample with a psychometrically refined approach to estimating individual ASC scores. Critically, this will allow us to test the potency of the ASC in also capturing life span trajectories and the ASC’s utility in explaining differences in predictive processing in language.

Previous studies have shown that across sensory modalities, older adults rely more heavily on prior expectations, a pattern that has been interpreted as a compensatory mechanism in response to age-related decline in executive functioning and sensory precision (14–16). We therefore first hypothesised that this shift might also manifest in changes in individual cognitive-perceptual style, with older adults potentially displaying a more prior-driven and thus more “schizotypal” mode of perceptual inference compared to younger individuals.

Second, we tested the functional relevance of these individual differences in a self-paced reading paradigm (16). Specifically, we asked whether variation along the ASC predicts behavioural sensitivity to lexical unpredictability (i.e., word surprisal), and whether this relationship is modulated by age. In doing so, we aimed to disentangle age-related effects from interindividual variation in cognitive-perceptual style, and to establish the ASC as a lifespan-sensitive dimension constraining predictive language processing.

## Results

We here report data from a large pooled sample comprising 340 participants (18-82 years; Mage = 38.94 ± 17.11 years; 59.12% female) who completed the AQ and SPQ questionnaires (10, 17) (analysis on full sample) as well as data from a smaller subset of the pooled sample, comprising 247 participants (Mage = 42.78 ± 17.35 years; 18–82 years, 55.87% female) who additionally performed a self-paced reading task (see Fig. 1b).

Individual positions on the Autism-Schizotypy Continuum (ASC), ranging from a more autism-like (low ASC scores) to a more schizotypal cognitive-perceptual style (high ASC scores), were captured by the second principal component of of AQ and SPQ subscale scores. Rather than using the original AQ scoring approach (17), we adopted a more recent method (18, 19) with improved psychometric properties (20).

Please refer to the *Methods* section for further details on the individual lines of analysis as well as to the *Supplementary Materials* for further details on how we computed individual ASC scores and validated our revised approach.

### Age shifts cognitive-perceptual style toward the autism pole

We first assessed how ASC scores change as a function of age, educational attainment, and gender using a linear model of the full pooled sample (N = 340, see age distribution in Fig. 2) that also controlled for a potential social desirability bias by controlling for the presence of an experimenter during the experiment.

**Figure 2.**
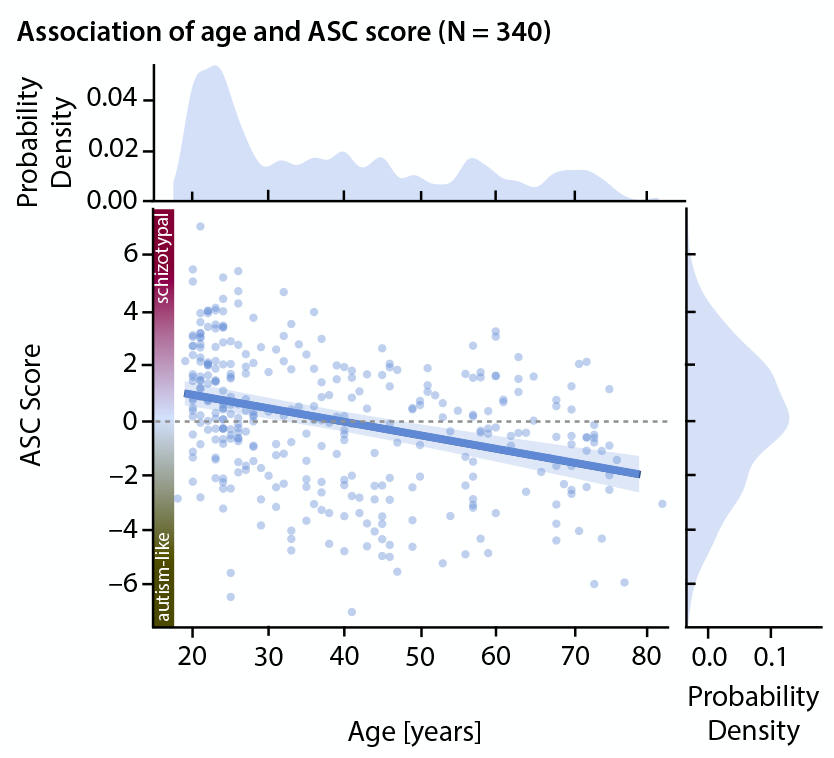
Age-related differences in cognitive-perceptual style (N = 340). Estimated marginal mean the effect of age on ASC scores, alongside the original data points (innermost panel) and marginal tributions of ASC-scores (left) and age (bottom). Older participants exhibited significantly lower ASC res – indicative of a more autism-like profiles – compared to younger participants.

Contrary to our hypothesis, we found that ASC scores decreased with age (*β* = –0.050, *t*(335.0) = –4.521, *p* < 0.001; see Fig. 2). We thus observe a continuous shift with age away from a schizotypy-like cognitive-perceptual style. Similarly, self-identified male participants exhibited lower scores than female participants (*β* = 1.001, *t*(335.0) = –4.031, *p* < 0.001; see Fig. S2; see Table S3 for full model details), suggesting gender-related differences in cognitive-perceptual style, with male individuals exhibiting more autism-like profiles than female individuals. No significant relationship was observed between educational attainment and individual ASC scores.

To evaluate the generalisability of our results, we employed 5-fold cross-validation (21). Across the five runs, the effects of age and gender consistently reached significance, suggesting that these effects are not dependent on specific subsets of the data (further details are provided in the *Supplementary Materials*; please see Fig. S2 and Table S6).

### Age and sensory-driven processing style amplify sensitivity to lexical surprisal

To examine how predictive processing during self-paced reading varies along the ASC, we fit a linear mixed model (LMM) of word-level reading times, including individual ASC scores alongside established demographic, lexical, and experiment-specific predictors.

Consistent with our previous findings (16), we found that participants read more slowly with increasing age (*β* = 0.006, *t*(239.40) = 7.825, *p* < 0.001) and word surprisal (*β* = 0.001, *t*(3333.63) = 7.504, *p* < 0.001), and that age amplified the effect of surprisal (*β* = 0.0002, *t*(161362.62) = 13.886, *p* < 0.001; see Fig. 3a). Put simply, surprisal effects increased in magnitude with advancing age – a pattern we discuss in more detail in our previous study (16).

**Figure 3.**
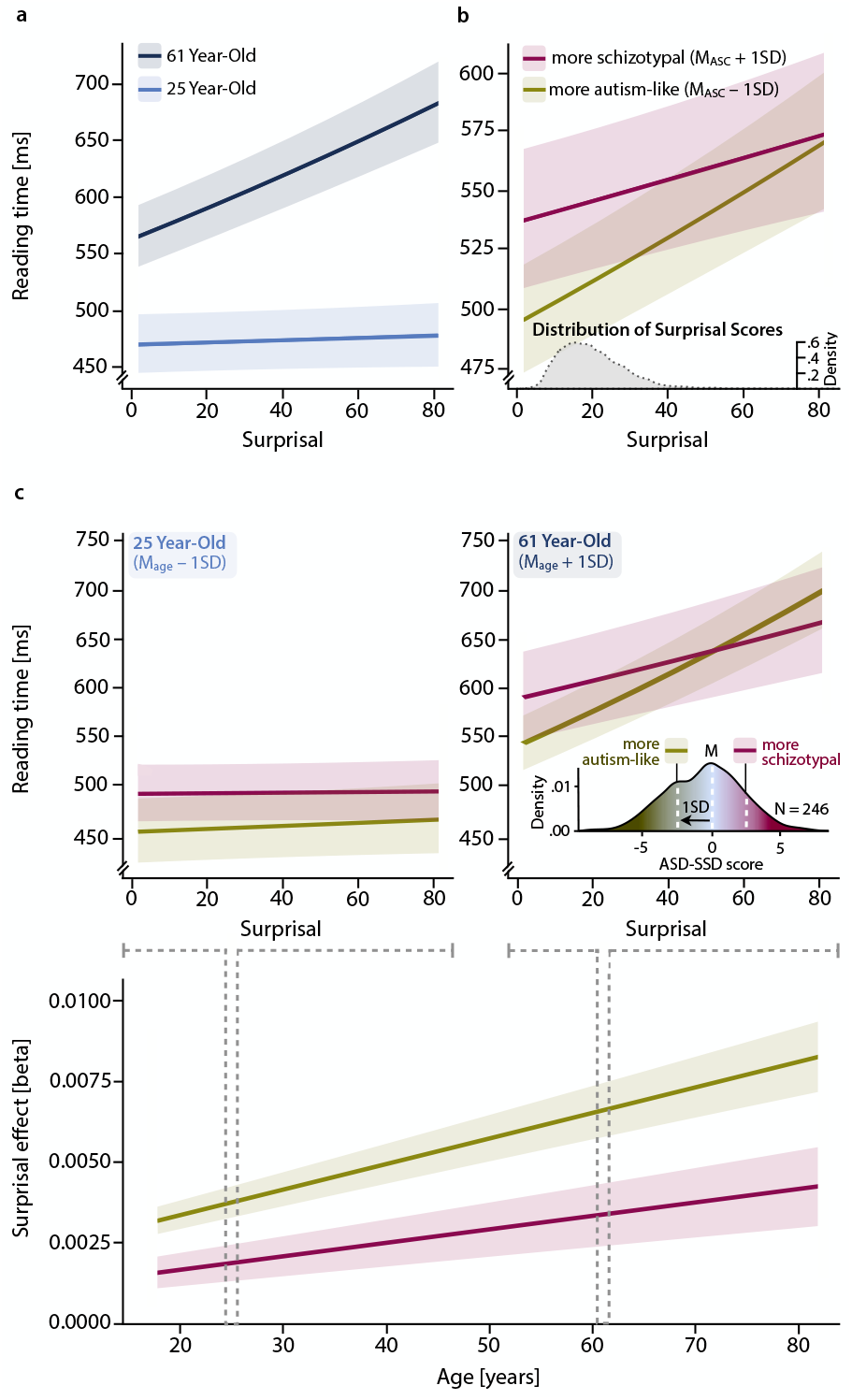
Age and cognitive-perceptual style jointly shape the response to unexpected lexical ut. **(a) Age increases sensitivity to unexpected input.** Panels show model-predicted marginal ect of surprisal on reading times for an exemplary younger (mean age −1 SD; 25 yr) and older (mean e +1 SD; 61 yr) individual. **(b) Autism-like profiles increase sensitivity to unexpected input**. nels show model-predicted marginal effect of surprisal in individuals with more autism-like (ASC −1) versus more schizotypal (ASC +1 SD) cognitive-perceptual styles. **(c) Age further amplifies the ect of surprisal in individuals with autism-like profiles**. Top panels show model-predicted rginal interaction effects of ASC score and age. Lower panel contrasts the magnitude of surprisal ects between autism-like and schizotypal styles directly. *Inset:* distribution of ASC scores (N = 247) d the M ± 1 SD cutoffs used for the modelled contrasts.

The novel contribution of this model lies in the inclusion of individual ASC scores as a predictor. Overall, higher ASC scores were associated with increased reading times (*β* = 0.013, *t*(239.32) = 2.284, *p* = 0.028), indicating that individuals with a more schizotypal processing style read more slowly than those with a more autism-like cognitive-perceptual style. Interestingly, higher word surprisal exerted a stronger effect on reading times in individuals with a more autism-like style (i.e., lower ASC scores; *β* = –0.0002, *t*(161141.90) = –5.999, *p* < 0.001; see Fig. 3b). This effect was further amplified by age: a significant three-way interaction between age, surprisal, and ASC score (*β* = –0.00001, *t*(160734.68) = –4.004, *p* < 0.001) revealed that older adults at the more autism-like end of the continuum were particularly sensitive to changes in word surprisal (see Fig. 3c and Table S7 for full results; see control analysis on the contribution of individual AQ and SPQ subscales to the 3-way interaction in the *Supplementary Materials*). Please note that the results reported here remain robust after excluding trials with extreme surprisal values (outside the M ± 2 SD range). Five-fold cross-validation (21) confirmed the robustness of the three-way interaction between surprisal, age, and ASC score. The effect was directionally consistent across all runs and reached significance in three of five models, supporting its stability and generalisability (see Fig. S3 and Table S8).

### Age-related shift toward autism pole of cognitive-perceptual style associated with poorer social functioning

In a last exploratory analysis, we aimed to get a more nuanced grasp of the age shift along the ASC. In a linear model, we predicted individual’s chronological age (log transformed) from their AQ and SPQ subscale scores. We found that older age was significantly associated with lower self-reported scores on several SPQ subscales, including *Ideas of Reference* (IR), *Social Anxiety* (SA), and *Unusual Perceptual Experiences* (UPE), as well as on the AQ subscale *Repetitive or Restricted Behaviour* (RRB). In contrast, older age was significantly associated with higher scores on the SPQ subscale *Odd Beliefs or Magical Thinking* (OB), as well as with AQ subscale scores reflecting difficulties in *Imagination* (I) and *Social Skills* (SS; see Table S5).

To relate these effects to the ASC construct, we considered subscale loadings on the ASC principal component. The SPQ subscales all exhibit relatively low absolute loadings on the ASC component (M = 0.242 ± 0.128), whereas AQ subscales show markedly higher loadings (M = 0.535 ± 0.282). Notably, the *Social Skills* subscale contributes disproportionately to the ASC score, with a particularly high loading towards the autism pole (loading: 0.706). This implies that changes in AQ subscales – especially *Social Skills* – exert a stronger influence on the ASC score than changes in SPQ subscales (see Table S4). Taken together, we found that age-related variation in ASC scores is strongly influenced by AQ subscales, particularly social skills, with additional age-related reductions in SPQ subscales such as *Unusual Perceptual Experiences* (loading: –0.455) and *Ideas of Reference* (–0.415) jointly contributing to an overall shift toward more autism-like profiles with age.

## Discussion

Healthy individuals demonstrably vary in cognitive-perceptual style along a continuum from more autism-like to more schizotypy-like profiles – the Autism-Schizotypy Continuum (ASC). Here, we asked how these individual differences in cognitive-perceptual style and its variation with age jointly shape predictive processing during naturalistic language comprehension.

First, our data reveal a systematic shift in cognitive-perceptual style towards less schizotypal profiles with age, driven by a decrease in several schizotypal subdimensions alongside increases in more trait-like aspects of social functioning.

Second, they show how the well-established increase in reading times with higher word-level surprisal (16, 22, 23) was systematically modulated by both age and cognitive-perceptual style affording novel insights into predictive processing: individuals with more autism-like profiles showed a stronger coupling between surprisal and reading times, and effect that was further amplified with age. We observed the strongest surprisal sensitivity in older individuals with more autism-like profiles. Notably, these effects remained robust under cross-validation.

These results are particularly striking given that differences in cognitive-perceptual style might be expected to emerge primarily under conditions of ambiguity or uncertainty (e.g., see 8, 24). Here, in contrast, the linguistic input used was highly predictable and fully intelligible. Nevertheless, ASC variation reliably and systematically modulated sensitivity to lexical surprisal even within this stable and strongly constrained processing environment, suggesting that the effects reported here may constitute a comparatively conservative estimate of cognitive-perceptual style differences in predictive processing.

Taken together, our data reveal a robust behavioural phenotype – increased sensitivity to contextually unexpected linguistic input – that varies systematically both across individuals and across the adult lifespan.

### Linking behavioural signatures to computational mechanisms

A main line of enquiry in our study was to employ individual ASC scores for understanding individuals’ predictive processing and surprisal sensitivity during self-paced reading. Such an effect would provide important evidence that inter-individual differences in cognitive-perceptual style along the ASC continuum indeed manifest behaviourally.

At first glance, the convergence of aging and autism-like profiles on stronger surprisal effects appears counterintuitive. Aging is often assumed to increase reliance on prior knowledge as a strategy to offset sensory decline (14, 16, 25), whereas autism-related accounts have been associated with weakened priors or enhanced weighting of sensory information(16)(26–29).

A key point, however, is that the same behavioural signature – here, stronger sensitivity to lexical surprisal – can arise from distinct computational mechanisms. In predictive processing terms, surprisal essentially corresponds to a form of prediction error (30, 31). The downstream behavioural consequence of such prediction errors depends on how strongly they influence ongoing inference and subsequent iterative belief updating. Critically, this influence can be modulated at different levels of the inferential hierarchy (32)

At a local level, they may reflect increased precision-weighting of incoming information, leading to a tighter coupling between prediction error and behaviour. Such an interpretation is consistent with accounts of autism-like traits emphasizing enhanced sensitivity to incoming sensory evidence (26–29). At a more global level, however, similar effects can emerge from differences in belief updating dynamics: if the environment is inferred to be volatile, priors are updated more rapidly and fail to stabilize, increasing the effective influence of prediction errors over time.

While these accounts converge behaviourally, they are computationally distinct, and the present stationary reading paradigm does not fully allow to adjudicate between them. Accordingly, our findings are best understood as reflecting trait- and age-dependent differences in the effective impact of prediction errors, potentially arising from multiple sources.

In the following, we consider in more detail how cognitive-perceptual style and age may give rise to this shared behavioural pattern via partially distinct mechanisms.

### Cognitive-perceptual style and belief updating during reading

Within this framework, the observed modulation of surprisal effects by an individual’s position along the ASC can be interpreted as reflecting differences in how incoming linguistic information updates current beliefs. Individuals with more autism-like profiles exhibited stronger surprisal effects, suggesting a tighter coupling between moment-to-moment input and predictive processing dynamics.

This pattern is consistent with predictive-processing accounts that attribute autism-like traits to increased precision-weighting of prediction errors or elevated estimates of environmental volatility (33, 34), both of which would increase reliance on incoming evidence. Conversely, individuals with more schizotypal profiles showed attenuated surprisal effects, consistent with a reduced impact of local prediction errors.

At a more abstract level, these differences in susceptibility to incoming input may manifest as variability in what could be described at the behavioural level as *uncertainty tolerance (33–35)*: Although the sensory input itself is constant across observers, its effective weighting may differ. Individuals assigning higher precision to incoming signals treat deviations as more informative, leading to stronger behavioural consequences of prediction errors, whereas individuals assigning lower precision exhibit a broader tolerance for ambiguity and a weaker impact of such deviations.

Importantly, this interpretation reframes the ASC not as indexing a simple trade-off between prior expectations and sensory evidence (6, 8, 12), but as capturing trait-like differences in belief updating regimes, that is, how strongly and how flexibly new information is incorporated into ongoing inference.

### Aging and the cost of prediction errors

The interaction between cognitive-perceptual style and word predictability, the central pattern in the present data, was strongest in older adults. Specifically, older individuals with a more autism-like profile exhibited the strongest modulation of reading times by word predictability, characterized by greater slowing for low-predictability words and greater facilitation for high-predictability words relative to their age-matched peers with a more schizotypal profile.

At a descriptive level, one possibility is that this interaction reflects a restricted dynamic range in younger readers, whose overall faster reading times may limit the observable expression of differences in predictive processing (36, 37). In this sense, age may partly modulate the visibility of the effect.

However, beyond such scaling considerations, aging likely affects the computational locus at which prediction-related effects arise. As outlined above, stronger surprisal effects need not necessarily reflect increased weighting or updating of prediction errors per se but may also arise from increased costs of resolving them. In the context of aging, unexpected words may place greater demands on lexical integration, revision of ongoing interpretations, or cognitive control processes, leading to amplified behavioural consequences of prediction errors (38).

Thus, while autism-like profiles may increase surprisal *sensitivity* by altering the influence of incoming information on belief updating, aging may primarily amplify the *cost* of processing prediction errors once they occur. The pronounced effects observed in older individuals with more autism-like profiles suggest that these mechanisms can combine, resulting in particularly strong behavioural signatures when both heightened sensitivity to input and increased integration costs are present.

### Age-related shifts in cognitive-perceptual style

To fully understand how cognitive-perceptual style shapes predictive processing across the lifespan, it is also important to consider how individual position along the ASC itself varies cross-sectionally with age. In the present data, older adults displayed a shift toward more autism-like profiles, an initially counterintuitive finding given the sensory decline typically associated with healthy aging (5, 39).

Closer inspection suggests that this shift is not driven by a uniform increase in autism-like traits, but rather by differential changes across subdimensions. Specifically, increasing age was associated with lower scores on subscales such as Ideas of *Reference, Social Anxiety, Unusual Perceptual Experiences*, and *Repetitive or Restricted Behaviour*, dimensions that capture responses to ambiguity, uncertainty, or perceived threat. This pattern is consistent with evidence that emotional reactivity to ambiguous or negatively valenced stimuli decreases with age, alongside improvements in emotional regulation (40–42).

At the same time, age was positively associated with higher scores on subscales such as *Odd Beliefs* or *Magical Thinking, (poor) Imagination*, and *(poor) Social Skills*, with changes in social functioning exerting a particularly strong influence on ASC scores. This suggests that while reactive or affective components of schizotypy may diminish with age, more stable aspects of cognitive-perceptual functioning, and their impact on social behaviour, may persist or even become more pronounced.

Taken together, these findings indicate that the observed age-related shift along the ASC may reflect, at least in part, changes in emotional reactivity and social functioning rather than a pure shift in perceptual inference mechanisms. This raises an important measurement issue: the ASC may conflate multiple dimensions that follow distinct developmental trajectories. Future work should therefore aim to more precisely isolate those components of the continuum that directly index cognitive-perceptual inference, for example by prioritizing subscales that capture belief formation and perceptual interpretation over those reflecting emotional responses or social functioning.

### Costs and benefits of predictive language processing

Finally, the pronounced susceptibility to lexical predictability in older individuals with more autism-like profiles provides a useful entry point into the ongoing debate surrounding the costs and benefits of predictive language processing (43). Previous work suggests that predictability facilitates processing when expectations are confirmed, but can incur costs when they are violated (43–45), although the extent and nature of these costs remain debated.

The present data indicate that both effects co-occur and vary systematically across individuals. Relative to older adults with a more schizotypal profile, who exhibited comparatively attenuated predictability effects, individuals with a more autism-like profile showed a stronger trade-off: faster processing for highly predictable words, but disproportionately slower processing when predictions were violated.

In naturalistic language, where highly predictable words are frequent (46), stronger engagement in predictive processing may therefore confer an overall advantage, despite occasional costs when expectations fail. Importantly, this trade-off is not fixed but varies across individuals and with age, underscoring that predictive processing is not uniformly beneficial but depends on the balance between expectation-driven facilitation and error-related costs.

## Conclusion

Our findings establish the Autism-Schizotypy Continuum (ASC) as a meaningful axis of inter-individual variability in predictive processing, indexing differences in belief updating rather than a simple balance between priors and sensory input. At the same time, the convergence of aging and autism-like profiles on stronger surprisal sensitivity suggests that similar behavioural signatures can arise from distinct computational routes, including differences in updating dynamics and the cost of resolving prediction errors. Together, these results argue for individualised hierarchical models of predictive processing that jointly account for individual traits and lifespan-related change.

## Materials and Methods

### Participants

The sample comprised 393 native German participants aged 18 to 82 years (M = 38.40 ± 16.78 SD years) with a balanced gender distribution of 53.69% male and 40.20% female participants. We pooled questionnaire data of six studies where participants were asked to complete the Autism Spectrum Quotient (AQ; 17) and the Schizotypal Personality Questionnaire (SPQ; 10). Participants completed the questionnaires in digital format either online (N = 233) or in a controlled lab session (N = 127), while a smaller subset (N = 33) completed them in pen-and-paper format at home.

Participants were recruited via convenience sampling, primarily through personal networks, through the online platform Prolific, and the participant databases of the University of Lübeck and the Max Planck Institute in Leipzig. All participants were native German speakers with no history of psychiatric or neurological disorders or substance abuse. Individuals who had consumed drugs or alcohol immediately prior to the measurement were not eligible for participation in the respective study and were therefore not included in this sample.

For further details on the subsamples used in the different analyses in the present study, please refer to the *Supplementary Materials*.

### Ethics statement

In all studies, participation was contingent upon providing written informed consent. Participants were compensated either financially (€12/h) or with course credits. All studies were conducted in accordance with the Declaration of Helsinki and were approved by the local ethics committees of the University of Lübeck and Leipzig University.

### ASC scores

#### Derivation of individual ASC scores

In all six studies, participants were asked to complete both the 50-item version of the AQ (17) and the 74-item version of the SPQ (10), which assess subclinical autistic and schizotypal traits, respectively. The SPQ uses a binary answer format, with all questions being endorsed “yes” scoring one point. They are all phrased in the same direction, such that a “yes” response consistently indicates more schizotypal cognition, perception, or behaviour; therefore, no reverse-keying is required. Items were scored and aggregated into the nine original subscales – ideas of reference (IR), Excessive Social Anxiety (SA), Odd Beliefs or Magical Thinking (OB), Unusual Perceptual Experiences (UPE), Odd or Eccentric Behaviour (OBH), No Close Friends (NCF), Odd Speech (OS), Constricted Affect (CA) and Suspiciousness (SUS) – following the original framework proposed by (10). For the AQ, subscale scores were derived using a revised five-subscale factor model as suggested by English et al. (20), instead of the original approach by Baron-Cohen et al. (17), which is usually employed when calculating ASC scores. The five-subscale system by Kloosterman et al. (18) comprises the subscales Social Skills (SS), Communication/Mindreading (CM), Restricted/Repetitive Behaviour (RRB), Imagination (I) and Attention to Detail (AD). A complete item-to-subscale mapping for the AQ is provided in the *Supplementary Materials* (see Table S2).

Prior to further analyses, we assessed the factorability of the correlation matrices underlying combinations of the SPQ subscale scores with the AQ subscale scores. For this purpose, we computed the *Kaiser-Meyer-Olkin Measure of Sampling Adequacy* (KMO-MSA; (47)) as well as *Bartlett’s Test of Sphericity* (48). Following these factorability checks, we conducted two *Principal Component Analyses* (PCA) to derive loadings for weighting AQ and SPQ subscales in the computation of individual ASC scores. When PCA is applied to AQ and SPQ subscale scores, it typically reveals two orthogonal components: While the first Principal Component (PC1) captures commonalities between the cognitive-perceptive styles of ASD and SSD, the loadings of the second Principal Component (PC2) distinguish between autism-like and positive schizotypal traits (7–9, 49). Accordingly, we extracted loadings from PC2 which we then multiplied with the *z*-scored AQ and SPQ subscale scores. This yielded one ASC score per participant, reflecting their position along the diametric axis defined by PC2. For further information on how we computed the ASC scores and how we validated this revised approach against two other models, please refer to the *Supplementary Materials*.

### Statistical analysis of ASC scores

To examine the potential influence of age, gender and education on individual cognitive-perceptual style, we fit a linear model with ASC scores as the dependent variable, with age, gender, and education as predictors. Additionally, as some participants completed the questionnaires in a controlled lab environment in the presence of at least one experimenter, we accounted for potential social desirability bias by including a binary predictor (‘presence of experimenter’). For this line of analyses, we employed a subsample of 340 participants. For some participants, only their highest academic degree was available, whereas years of education were missing. In these cases, missing values were approximated using the minimum standard duration typically required to obtain the reported degree. This approximation did not account for individual variations such as skipping years or alternative educational trajectories. All continuous predictors were mean-centred prior to analysis. The full model formula was:

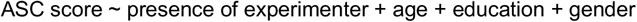

To further assess the robustness of our effects, we employed 5-fold cross-validation (21). Specifically, we divided the full sample of 340 participants into five folds of 68 participants, stratified by gender. The resulting folds – and the original full sample – were highly comparable in their distributions of gender, age, and education (see Fig. 2c). This ensured that each fold provided a balanced representation across the key demographic variables and adequately covered the full range of combinations of age, education, and gender.

### Self-paced reading task

#### Experimental design

A subsample of 247 participants completed a self-paced reading task prior to questionnaire completion (see Fig. 1b). Participants read short newspaper articles (300 words each) word-by-word at their own pace, advancing through the text by pressing the space bar. Following each article, they answered three multiple-choice comprehension questions. To manipulate cognitive load, some blocks included an orthogonal n-back task based on the font colour of the words, which participants practiced separately at the start of the experiment (see (16) for further details on the task). Please note that the present study focused on the *Reading Only* (self-paced reading without the n-back) condition to characterize baseline reading behaviour without cognitive load interference.

Following our previous study (16), we used word-level reading times alongside comprehension accuracy calculated at both the participant and block levels as the percentage of correctly answered questions. Furthermore, we derived a mean d-prime score for each participant, indexing working memory capacity, from the non-verbal single-task n-back training blocks.

### Generation of word surprisal and entropy scores

For all texts presented in the reading task, we computed word-level surprisal – the negative logarithm of a word’s probability given its preceding context (31) – as a measure of how unexpected a word is. Higher surprisal values therefore indicate greater unpredictability and increased processing difficulty upon encountering the respective word (30, 31, 50).

Additionally, we extracted word entropy, defined as the mean surprisal of all possible continuations in the vocabulary (51, 52). Entropy thus captures the uncertainty about which word will occur next prior to its presentation – that is, the unpredictability inherent in the prediction itself. Together, surprisal and entropy thus reflect complementary aspects of word predictability: entropy indexes pre-stimulus uncertainty, whereas surprisal reflects post-stimulus processing difficulty. In the present study, word surprisal is used to operationalise word (un)predictability, while entropy is included solely as a lexical control variable in the models.

Surprisal and entropy were computed using a 12-layer GPT-2 model (53) pre-trained on a corpus of German texts by the MDZ Digital Library team (dbmdz) at the Bavarian State Library, along with the corresponding tokenizer, both obtained from the Hugging Face model hub (54). Following our previous study (16), we employed a context window comprising the two preceding words. All computations were performed in Python version 3.10.12 (55).

### Preprocessing and statistical analysis of reading times

To examine how individual cognitive-perceptual style relates to participants’ perceptual processing, we analysed self-paced reading times. To this end, we excluded all blocks in which participants failed to answer any comprehension questions correctly, and removed outlier trials from the remaining blocks based on excessively short or long reading times.

To assess the joint influence of age, cognitive-perceptual style, and word surprisal on reading at the single-word level, we fit a linear mixed-effects model (LMM) on log-transformed reading times. Model specification closely followed our previous study (16) and encompassed task- and experiment-related control variables alongside the predictors of interest. Random effects included random intercepts for participants as well as the effect of text and the current word. All continuous predictors were mean-centred. For further details on outlier exclusion and model specifications, please refer to the *Supplementary Materials*.

To probe the robustness and generalisability of the three-way interaction of age, surprisal and ASC scores, we employed 5-fold cross-validation (21), employing folds consisting of 49 or 50 participants each.

log(RT) ~ RT of previous word +

> mean d-prime from single tasks +
>
> mean comprehension question performance +
>
> block-level deviation from mean comprehension question performance +
>
> recording location + word entropy +
>
> word frequency + word length (without punctuation) + block number + trial number +
>
> **surprisal * age * ASC score** +
>
> (1 | ID) + (1 | text number) + (1 | word)

*Note. Model structure. RT = Reading Time, ID = participant*.

The experiment was implemented in *lab*.*js* (56), and either hosted online on *OpenLab* (57) with data being saved on *OSF* (58), or run locally as controlled lab experiments. All data analyses were performed in *R* (version 4.4.1, 59) using R’s packages *factoextra* (60), *psych* (61), *lme4* (62), *lmerTest* (63) and *sjPlot* (64)).

## Acknowledgments

We thank Luca Tarasi for his generous advice on computing ASC scores and for kindly sharing his data with us.

## Declaration of AI and AI-assisted technologies in the writing process

During the preparation of this work the authors used GPT-4o-mini (OpenAI, Inc.) to rephrase sentences. The authors reviewed and edited any AI-generated content as needed and take full responsibility for the content of the publication.

## Data availability

All experimental and analysis scripts as well as preprocessed data are publicly accessible on OSF (Project Link: https://osf.io/8scnf).

## Author Contributions

M.S.: Conceptualization, data curation, formal analysis, investigation, methodology, project administration, visualisation, writing: original draft, review & editing

S.T.: Formal analysis, methodology, writing: review & editing

S.M.: Methodology, writing: review & editing

G.H.: Methodology, funding acquisition, supervision, writing: review & editing

J.O.: Conceptualization, formal analysis, methodology, funding acquisition, supervision, writing: review & editing

