## Supplements for "Cognitive-Perceptual Style on the Autism-Schizotypy Continuum Shapes Surprisal Sensitivity"

### Supplementary methods

#### *Sample characteristics*

Some analyses in the present study were conducted on subsamples of the full dataset, depending on data availability and task completion. To compute the factor loadings used to derive ASC scores, we employed the full sample of 393 participants.

For examining the effects of age, education, and gender on ASC scores, we included all participants with complete information on these variables. We excluded individuals with missing data and those reporting implausibly low education values (e.g., less than five years of schooling). This resulted in a subsample of *N* = 340 participants aged 18 to 82 years (M = 38.94 ± 17.11 years; 59.12% female), with a mean of 14.21 ± 2.41 years of schooling.

For investigating the relationship of individual scores on the ASC with behavioural markers of predictive processing, we employed a sub-sample of N = 247 participants aged 18 to 82 years (M = 42.78 ± 17.35 years; 55.87% female) who had performed a self-paced reading task in addition to completing the questionnaires.

As reading time data were pooled from three related experiments, minor procedural variations existed: participants completed either two or three Reading Only blocks (two blocks: N = 185, sub-sample from Schuckart et al. [2024]; three blocks: N = 62 from a new sample), resulting in some participants contributing more data than others. Texts were randomly selected from a pool of ten different texts. Two of the studies used *lab.js* (56) for implementation, with experiments run either locally in a controlled lab setting or online via *OpenLab* and *OSF* for hosting and data storage (57, 58). The third study employed Python with the *PsychoPy* package (65) and was run locally in a controlled lab setting.

#### *Derivation of individual ASC scores*

In all six studies, participants were asked to complete both the 50-item version of the AQ (17) and the 74-item version of the SPQ (10).

The SPQ uses a binary answer format, with all questions being endorsed "yes" scoring one point. Items were scored and aggregated into the nine original subscales, following the original framework proposed by (10).

For the AQ, subscale scores were derived using three different methods: First, we applied the five "classic" subscales proposed by (17) and used in the studies by Tarasi et al. (7, 8, 66, 67), which reflect poor Social Skills (SS), Attention-Switching (AS), Communication Skills (CS), and Imagination (I) as well as exceptional Attention to Detail (AD). In the AQ, we applied the classical scoring method in which the 4-point Likert scale is collapsed into a binary format: responses indicating either mild or strong endorsement of autism-like behaviour, perception, or cognition are scored as 1, and all others as 0. This approach reflects the traditional scoring procedure (17), despite the underlying Likert response format.

Second, we also computed AQ subscale scores using two more recent, alternative factor models as suggested by (20): the three-subscale system by (68), which includes the subscales Social Skills (SS), Details/Patterns (DP), and Communication/Mindreading (CM); and the five-subscale system by (18), comprising Social Skills (SS), Communication/Mindreading (CM), Restricted/Repetitive Behaviour (RRB), Imagination (I) and Attention to Detail (AD). A complete item-to-subscale mapping is provided in Table S2.

Prior to further analyses, we assessed the factorability of the correlation matrices underlying combinations of SPQ subscale scores with AQ subscale score derived from three different factor models: Baron-Cohen, Russell-Smith, and Kloosterman. For this purpose, we computed the *Kaiser-Meyer-Olkin Measure of Sampling Adequacy* (KMO-MSA; (47)) as well as *Bartlett's Test of Sphericity* (48).

As KMO-MSA results indicated acceptable factorability only for the Baron-Cohen and Kloosterman models, we limited subsequent analyses to comparisons based on the Baron-Cohen- and Kloosterman-based AQ subscales and their corresponding ASC scores.

Following these factorability checks, we conducted two *Principal Component Analyses* (PCA) to derive loadings for weighting AQ and SPQ subscales in the computation of individual ASC scores. One PCA included all SPQ subscales along with the Baron-Cohen-based AQ subscales; the other combined SPQ subscales with the Kloosterman-based AQ subscales (see Fig. S1). In line with previous studies deriving individual ASC scores using PCA, we used unrotated PCAs in the present analyses.

When PCA is applied to AQ and SPQ subscale scores, it typically reveals two orthogonal components: While the first Principal Component (PC1) captures commonalities between the cognitive-perceptive styles of ASD and SSD, the loadings of the second Principal Component

(PC2) distinguish between autistic-like and positive schizotypal traits (7–9, 49). Accordingly, we extracted loadings from PC2 to compute individual ASC scores.

It is important to note that our reported loadings show a sign flip relative to those reported by (8). This is a known and purely arbitrary feature of PCA: As PCA is based on eigenvectors, the direction (sign) of a component is mathematically undefined – flipping all signs yields an equivalent solution (69). As a result, the sign of each component can differ across samples or software implementations without affecting the underlying structure. However, to facilitate interpretability and ensure comparability across studies, we subsequently aligned the direction of PC2 with previous work by flipping all signs such that higher ASC scores reflect more schizotypal traits and lower scores reflect more autism-like traits.

Employing the extracted loadings of PC2 from two PCAs, and the previously published loadings from (8), we computed individual ASC scores based on three distinct models, summarised in Table S1. Models 1 and 2 both use the SPQ item scoring and subscale structure from (10) as well as the item scoring and factor structure proposed by (17), differing only in the sample used to compute loadings (Tarasi's smaller, younger sample versus our larger, age-diverse sample). Model 3 applies the SPQ item scoring and factor structure from (10), combined with the more nuanced AQ item scoring from (19) and the alternative, more psychometrically sound factor structure proposed by (18), also computed using questionnaire data from our larger sample. This approach enabled us to validate our revised method (model 3) against both the original method as implemented here (model 2) and a previously published model of the same structure as our model 2, known to yield robust results in a psychophysical task (model 1).

For each model, we multiplied each PC2 loading vector with the respective *z*-scored subscale scores. This yielded one ASC score per participant for each model, reflecting their position along the diametric axis defined by PC2. We ran Spearman rank correlations between the ASC scores from each model to assess the consistency of the scoring approaches.

As we observed strong correlations between established ASC scores (models 1 and 2) and the revised scores (model 3), all subsequent analyses employed ASC scores computed with the revised scoring method.

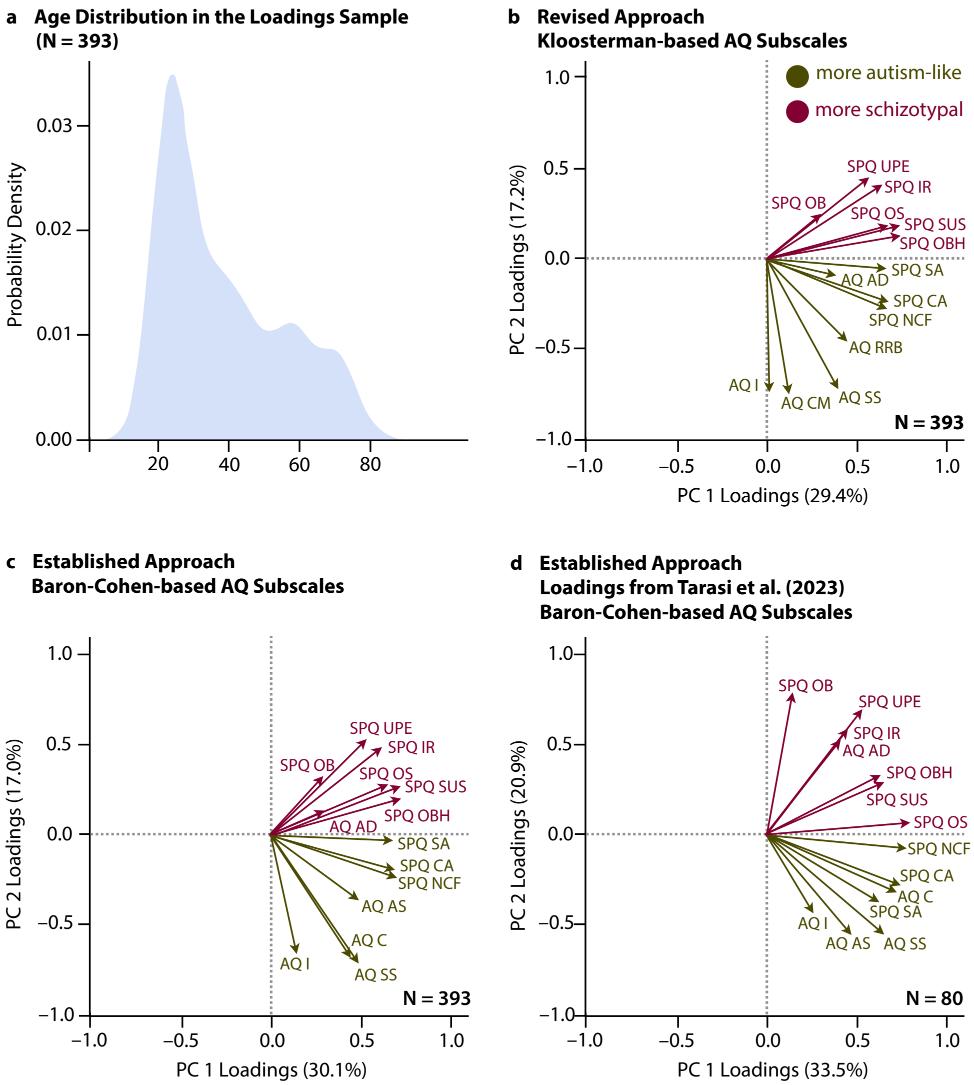


**Figure S1. Results of the Principal Component Analyses.** Panel **a** depicts the age distribution of the sample used to compute PCA loadings. Panel **b** illustrates the loadings for the first two principal components from the PCA on our revised scoring approach, which employed the AQ factor model proposed by Kloosterman et al. (18). In contrast, panels **c** and **d** display PCA results from the original scoring method, which relies on the Baron-Cohen five-subscale AQ model (17). The loadings in panels **b** and **c** are based on data from our large, age-diverse dataset (N = 393), while panel **d** reproduces the loadings reported by Tarasi et al. (8), which were computed with data from a smaller sample of young adults (N = 80). For clearer comparison, the loadings in panels **b** and **c** have been sign-flipped to align with those from Tarasi et al. An overview of the abbreviations for the AQ and SPQ subscales is provided in Table S4.

**Table S1.** Models for computing individual ASC scores.

| Model | SPQ item scoring and factor structure | AQ item scoring | AQ factor structure | Sample for computation of loadings |
|---|---|---|---|---|
| 1 | Raine (1991) | Baron-Cohen et al. (2001) | Baron-Cohen et al. (2001) | N = 80, young<br>from Tarasi et al (2023) |
| 2 | Raine (1991) | Baron-Cohen et al. (2001) | Baron-Cohen et al. (2001) | N = 393, age-diverse |
| 3 | Raine (1991) | Austin (2005) | Kloosterman et al. (2011) | N = 393, age-diverse |

**Table S2.** AQ items (17) alongside the subscales to which they are assigned in the factor structures proposed by Baron-Cohen et al. (2001), Russel-Smith et al. (2011), and Kloosterman et al. (2011).

| Item Nr | Subscale | | |
|---|---|---|---|
| | **Baron-Cohen et al. (2001)** | **Russel-Smith et al. (2011)** | **Kloosterman et al. (2011)** |
| 1* | Social Skills (SS) | Social Skills (SS) | Social Skills (SS) |
| 2 | Attention Switching (AS) | | Restricted / Repetitive Behaviour (RRB) |
| 3* | Imagination (I) | | Imagination (I) |
| 4 | Attention Switching (AS) | | Restricted / Repetitive Behaviour (RRB) |
| 5 | Attention to Detail (AD) | Details/Patterns (DP) | Attention to Detail (AD) |
| 6 | Attention to Detail (AD) | Details/Patterns (DP) | Attention to Detail (AD) |
| 7 | Communication (C) | | |
| 8* | Imagination (I) | | Imagination (I) |
| 9 | Attention to Detail (AD) | Details/Patterns (DP) | |
| 10* | Attention Switching (AS) | Social Skills (SS) | Communication / Mindreading (CM) |
| 11* | Social Skills (SS) | Social Skills (SS) | Social Skills (SS) |
| 12 | Attention to Detail (AD) | Details/Patterns (DP) | Attention to Detail (AD) |
| 13 | Social Skills (SS) | Social Skills (SS) | |
| 14* | Imagination (I) | | |
| 15 | Social Skills (SS) | Social Skills (SS) | Social Skills (SS) |
| 16 | Attention Switching (AS) | | |
| 17* | Communication (C) | Social Skills (SS) | Social Skills (SS) |
| 18 | Communication (C) | | Restricted / Repetitive Behaviour (RRB) |
| 19 | Attention to Detail (AD) | Details/Patterns (DP) | Attention to Detail (AD) |
| 20 | Imagination (I) | Communication / Mindreading (CM) | Imagination (I) |
| 21 | Imagination (I) | | Imagination (I) |
| 22 | Social Skills (SS) | Social Skills (SS) | Social Skills (SS) |
| 23 | Attention to Detail (AD) | Details/Patterns (DP) | Attention to Detail (AD) |

| | | | |
|---|---|---|---|
| 24* | Imagination (I) | | |
| 25* | Attention Switching (AS) | | Restricted / Repetitive Behaviour (RRB) |
| 26 | Communication (C) | Social Skills (SS) | |
| 27* | Communication (C) | Communication / Mindreading (CM) | Communication / Mindreading (CM) |
| 28* | Attention to Detail (AD) | | |
| 29* | Attention to Detail (AD) | | |
| 30* | Attention to Detail (AD) | | |
| 31* | Communication (C) | Communication / Mindreading (CM) | Communication / Mindreading (CM) |
| 32* | Attention Switching (AS) | | |
| 33 | Communication (C) | | |
| 34* | Attention Switching (AS) | Social Skills (SS) | |
| 35 | Communication | Communication / Mindreading | |
| 36* | Social Skills (SS) | Communication / Mindreading (CM) | Communication / Mindreading (CM) |
| 37* | Attention Switching (AS) | | |
| 38* | Communication (C) | Social Skills (SS) | Social Skills (SS) |
| 39 | Communication (C) | Communication / Mindreading (CM) | Restricted / Repetitive Behaviour (RRB) |
| 40* | Imagination (I) | | Imagination (I) |
| 41 | Imagination (I) | Details/Patterns (DP) | |
| 42 | Imagination (I) | | |
| 43 | Attention Switching (AS) | | |
| 44* | Social Skills (SS) | Social Skills (SS) | Social Skills (SS) |
| 45 | Social Skills (SS) | Communication / Mindreading (CM) | Communication / Mindreading (CM) |
| 46 | Attention Switching (AS) | Social Skills (SS) | |
| 47* | Social Skills (SS) | Social Skills (SS) | Social Skills (SS) |
| 48* | Social Skills (SS) | Communication / Mindreading (CM) | |
| 49* | Attention to Detail (AD) | | |
| 50* | Imagination (I) | | |

#### *Analysis of reading times: Outlier exclusion criteria and specifications of the linear mixed-effects model*

To remove extended breaks from the reading time data, we first excluded pauses of 5000 ms or longer, which affected 0.8% of trials. We then applied a POMS transformation (70) to the reading times, followed by a $z$-transformation of the square-root–transformed values to identify and exclude statistical outliers (71, 72). On average, 2.05% of trials were excluded, with 72.724 ± 11.665 excluded trials per participant.

The linear mixed-effects model (LMM) fit on log-transformed reading times closely followed our previous study (16), incorporating the following continuous fixed effects: word frequency (as estimated using Python's *wordfreq* package; (73)), word length, and entropy as lexical measures; comprehension accuracy at the block and participant level as task performance measures; mean d-prime from a non-linguistic n-back task as a proxy for individual working memory capacity; log-transformed reading time from the previous word to account for potential sequential modulation effects if the previous trial was ended prematurely or the participant paused reading within a block; and finally block and trial number to control for training and fatigue effects. Recording location (lab vs. online) was included as a categorical control variable to account for potential differences in task engagement or familiarity with the used equipment.

Additionally, similar to the original model used in our previous study (16), we added age and word surprisal as continuous predictors to the model, alongside each participant's individual ASC score, as well as their two- and three-way interactions.

### Supplementary results

#### *Factorability checks*

Bartlett's test was significant for all three combinations of SPQ and AQ subscales (SPQ and Baron-Cohen-based AQ: $\chi^2(91) = 1986.059$, $p < .001$; SPQ and Russel-Smith-based AQ: $\chi^2(66) = 1618.067$, $p < .001$; SPQ and Kloosterman-based AQ: $\chi^2(91) = 1883.072$ $p < .001$), indicating that the subscales showed sufficient intercorrelation to proceed with principal component analysis.

However, the KMO-MSA results revealed insufficient sampling adequacy for the Russel-Smith based AQ subscales: the three AQ subscales yielded individual MSAs of 0.60 (RS-SS), 0.77 (RS-DP), and 0.46 (RS-CM), with one subscale falling below the commonly accepted threshold of 0.60 (47), suggesting limited shared variance with the remaining subscales. In contrast, sampling adequacy was acceptable for the other two models. For the Baron-Cohen-based AQ subscales, individual MSAs ranged from 0.65 to 0.82, with an overall MSA of 0.78. For the Kloosterman model, AQ subscale MSAs ranged from 0.72 to 0.80, with an overall MSA of 0.80.

Across all three models, SPQ subscales consistently showed good sampling adequacy, with individual MSAs averaging 0.807 ± 0.057. The lowest SPQ MSA was found for the subscale Odd Beliefs or Magical Thinking (OB; range of MSAs: 0.68–0.69), and the highest one for Odd or Eccentric Behaviour (OBH; range of MSAs: 0.87–0.89).

Taken together, as KMO-MSA results indicated acceptable factorability only for the Baron-Cohen and Kloosterman models, we limited all subsequent analyses to comparisons based on the Baron-Cohen- and Kloosterman-based AQ subscales and their corresponding ASC scores.

#### *Control analysis: 5-fold cross-validation of the demographic effects*

To evaluate the generalisability of our results, we employed 5-fold cross-validation (21). Across the five runs, the effects of age and gender consistently reached significance, suggesting that these effects are not overly dependent on specific subsets of the data (see Fig. S2 and Table S6). The average root mean squared error (RMSE) was 2.177 ± 0.305, indicating that, on average, predicted ASC scores deviated from observed scores by approximately 2.18 points. Considering that the linear model included only basic demographic predictors (age, education, and gender), this level of prediction error is reasonable. The model's explanatory power, as reflected in an average $R^2$ of 0.231 ± 0.014, further indicates that these variables capture meaningful variance in ASC scores despite the model's simplicity.

Across runs, we observed robustly lower ASC scores in self-identified men compared to women, which most likely arises from well-documented gender biases in the AQ and SPQ. As such, this observed gender effect should be interpreted cautiously, as it may reflect both genuine differences and measurement-related artefacts. Men typically score higher on the AQ subscales *social skill*, *attention switching*, *communication*, and *imagination* (17, 74, 75), whereas in schizotypy measures they score lower than women on perceptual-cognitive traits, but higher on social subscales (76–79). Since perceptual-cognitive schizotypy loads opposite to autistic traits on the ASC, while social traits load in the same direction, these patterns may bias composite ASC scores.

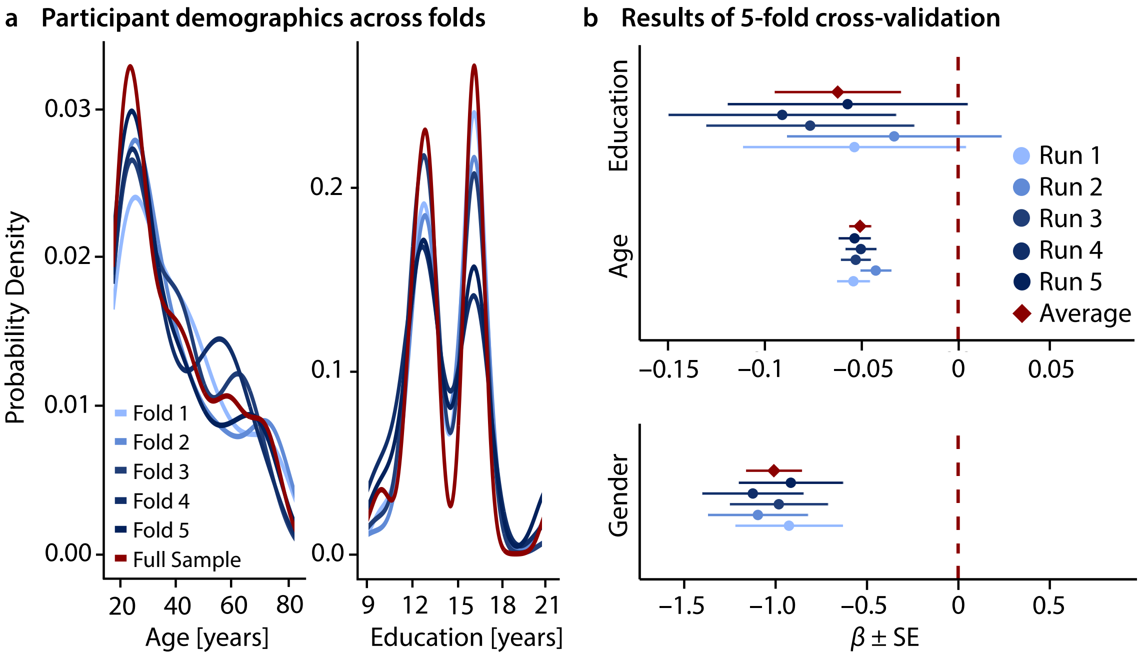


**Figure S2. Results of the 5-fold cross-validation (N = 340).** Panel **a** shows the distributions of age and education in all folds as well as in the original full sample, while panel **b** summarizes the cross-validation results, presenting estimates ± standard errors (SE). Average estimates were calculated across runs. To pool SEs, we averaged the squared SEs (variances) from each fold, added the variance of the estimates across folds adjusted for sample size, and then took the square root to reflect total uncertainty.

**Table S3.** Results for LM for effects of age, education, gender and the presence of an experimenter on individual ASC scores.

| ***Predictor*** | ***Estimate*** | ***Std. Error*** | ***CI*** | ***t*** | ***p*** | |
|---|---|---|---|---|---|---|
| Presence of experimenter | 1.137837 | 0.275175 | 0.59655 – 1.67913 | 4.134957 | $1.123 \times 10^{-4}$ | * |
| age | –0.049552 | 0.006952 | –0.06323 – –0.03588 | –7.127959 | $3.148 \times 10^{-11}$ | * |
| education | –0.060111 | 0.049877 | –0.15822 – 0.03800 | –1.205180 | $2.862 \times 10^{-1}$ | |
| gender | –1.000972 | 0.248288 | –1.48937 – –0.51257 | –4.031492 | $1.144 \times 10^{-4}$ | * |
| N | 340 | | | | | |
| df | 335.0 | | | | | |
| R2 / R2 adjusted | 0.230 / 0.221 | | | | | |

*Note*. All continuous predictors were mean-centred. Degrees of freedom for p-values and confidence intervals (CI) were computed using Wald's approximation. All p-values reported here are FDR-corrected and were computed using ANOVAs with type III sum of squares. Results that are significant on an alpha level of 0.05 are marked with a star.

**Table S4.** Loadings for all AQ and SPQ subscales.

| PCA Configuration | Subscales | Loadings on PC2 | |
|---|---|---|---|
| | | Sample from Current Study (N = 393) | Sample from Tarasi et al. (2023; N = 80) |
| SPQ and Baron-Cohen AQ Subscales | AQ: Social Skills (SS) | 0.698 | -0.54 |
| | AQ: Attention Switching (AS) | 0.354 | -0.54 |
| | AQ: Attention to Detail (AD) | -0.131 | 0.52 |
| | AQ: Communication (C) | 0.671 | -0.31 |
| | AQ: Imagination (I) | 0.645 | -0.42 |
| | SPQ: Ideas of Reference (IR) | -0.477 | 0.58 |
| | SPQ: Social Anxiety (SA) | 0.030 | -0.36 |
| | SPQ: Odd Beliefs / Magical Thinking (OB) | -0.318 | 0.78 |
| | SPQ: Unusual Perceptual Experiences (UPE) | -0.523 | 0.69 |
| | SPQ: Odd/Eccentric Behaviour (OBH) | -0.196 | 0.33 |
| | SPQ: No Close Friends (NCF) | 0.231 | -0.07 |
| | SPQ: Odd Speech (OS) | -0.276 | 0.07 |
| | SPQ: Constricted Affect (CA) | 0.190 | -0.27 |
| | SPQ: Suspiciousness (SUS) | -0.266 | 0.29 |

| PCA Configuration | Subscales | Loadings on PC2 | |
|---|---|---|---|
| | | Sample from Current Study (N = 393) | Sample from Tarasi et al. (2023; N = 80) |
| SPQ and Kloosterman AQ Subscales | AQ: Social Skills (SS) | 0.706 | |
| | AQ: Communication / Mindreading (CM) | 0.733 | |
| | AQ: Restricted / Repetitive Behaviour (RRB) | 0.438 | |
| | AQ: Imagination (I) | 0.718 | |
| | AQ: Attention to Detail (AD) | 0.080 | |
| | SPQ: Ideas of Reference (IR) | -0.415 | |
| | SPQ: Social Anxiety (SA) | 0.045 | |
| | SPQ: Odd Beliefs / Magical Thinking (OB) | -0.251 | |
| | SPQ: Unusual Perceptual Experiences (UPE) | -0.455 | |
| | SPQ: Odd/Eccentric Behaviour (OBH) | -0.133 | |
| | SPQ: No Close Friends (NCF) | 0.265 | |
| | SPQ: Odd Speech (OS) | -0.193 | |
| | SPQ: Constricted Affect (CA) | 0.229 | |
| | SPQ: Suspiciousness (SUS) | -0.193 | |

*Note.* Please note our loadings are sign-flipped compared to the ones from (8); all ASC subscale scores were therefore multiplied by –1 to align with the direction of the ASC reported in the literature.

### Table S5. Effect of AQ and SPQ subscales on age.

| ***Predictor*** | ***Estimate*** | ***Std. Error*** | ***CI*** | ***t*** | ***p*** | |
|---|---|---|---|---|---|---|
| AQ AD (exceptional Attention to Detail) | 0.010516 | 0.007934 | -0.005092 – 0.026125 | 1.325446 | $2.389x10^{-1}$ | |
| AQ CM (poor Communication/Mindreading) | 0.011530 | 0.008802 | -0.005785 – 0.028844 | 1.309969 | $2.389x10^{-1}$ | |
| AQ I (poor Imagination) | 0.029906 | 0.008306 | 0.013567 – 0.046245 | 3.600662 | **$2.749x10^{-3}$** | * |
| AQ SS (poor Social Skills) | 0.022391 | 0.006422 | 0.009757 – 0.035025 | 3.486574 | **$2.779x10^{-3}$** | * |
| AQ RRB (Repetitive/Restricted Behaviour) | -0.024728 | 0.009537 | -0.043490 – -0.005965 | -2.592691 | **$3.135x10^{-2}$** | * |
| SPQ IR (Ideas of Reference) | -0.033796 | 0.014352 | -0.062028 – -0.005563 | -2.354841 | **$4.719x10^{-2}$** | * |
| SPQ SA (Social Anxiety) | -0.026523 | 0.011527 | -0.049200 – -0.003847 | -2.300927 | **$4.719x10^{-2}$** | * |
| SPQ OB (Odd Beliefs / Magical Thinking) | 0.046600 | 0.018095 | 0.011004 – 0.082197 | 2.575352 | **$3.135x10^{-2}$** | * |
| SPQ OBH (Odd / Eccentric Behaviour) | -0.013583 | 0.014686 | -0.042474 – 0.015308 | -0.924868 | $3.811x10^{-1}$ | |
| SPQ NCF (No Close Friends) | 0.014211 | 0.014013 | -0.013356 – 0.041777 | 1.014117 | $3.592x10^{-1}$ | |
| SPQ SUS (Suspiciousness) | 0.033695 | 0.018619 | -0.002932 – 0.070322 | 1.809771 | $1.069x10^{-1}$ | |
| SPQ CA (Constricted Affect) | -0.031024 | 0.015816 | -0.062138 – 0.000090 | -1.961509 | $8.444x10^{-2}$ | |
| SPQ OS (Odd Speech) | 0.002981 | 0.011187 | -0.019026 – 0.024988 | 0.266487 | $7.900x10^{-1}$ | |
| SPQ UPE (Unusual Perceptual Experiences) | -0.040471 | 0.018157 | -0.076191 – -0.004752 | -2.228910 | **$4.968x10^{-2}$** | * |
| N | 340 | | | | | |
| df | 328.0 | | | | | |
| $R^2$ / $R^2$ adjusted | 0.273 / 0.241 | | | | | |

*Note.* All continuous predictors were mean-centred. The dependent variable age was log-transformed prior to fitting. Degrees of freedom for *p*-values, standard errors and confidence intervals (CI) were computed using Wald's approximation. All *p*-values reported here are FDR-corrected and were computed using ANOVAs with type III sum of squares. Results that are significant on an alpha-level of 0.05 are marked with an asterisk. Please note AQ subscales are based on the Kloosterman et al. (2011) factor structure. Subscale scores were not weighted by loadings for this analysis.

**Table S6.** Results of the 5-fold cross-validation for the effects of age, gender, education and their respective interactions on ASC scores.

| Predictor | | run 1 ($N_{test}$ = 68, $N_{training}$ = 272) | run 2 ($N_{test}$ = 68, $N_{training}$ = 272) | run 3 (N = 69, $N_{training}$ = 271) | run 4 (N = 67, $N_{training}$ = 273) | run 5 (N = 68, $N_{training}$ = 272) |
|---|---|---|---|---|---|---|
| **experimenter present** | ***β*** | 1.259084 | 1.069518 | 1.170429 | 1.116494 | 1.057136 |
| | **Std. Error** | 0.3219330 | 0.3012710 | 0.2962803 | 0.3065856 | 0.3141862 |
| | **95% CI** | 0.625 – 1.893 | 0.476 – 1.663 | 0.587 – 1.754 | 0.513 – 1.720 | 0.439 – 1.676 |
| | ***t*** | 3.911013 | 3.550020 | 3.950410 | 3.641705 | 3.364680 |
| | ***p*** | **$1.166 \times 10^{-4}$** | **$4.551 \times 10^{-4}$** | **$1 \times 10^{-4}$** | **$3.250 \times 10^{-4}$** | **$8.788 \times 10^{-4}$** |
| **age** | ***β*** | -0.05277753 | -0.04151425 | -0.05161826 | -0.04903529 | -0.05218991 |
| | **Std. Error** | 0.008178704 | 0.007675585 | 0.007423284 | 0.007651037 | 0.007974477 |
| | **95% CI** | -0.069 – -0.037 | -0.057 – -0.026 | -0.066 – -0.037 | -0.064 – -0.034 | -0.068 – -0.036 |
| | ***t*** | -6.453043 | -5.408610 | -6.953561 | -6.408974 | -6.544618 |
| | ***p*** | **$5.134 \times 10^{-10}$** | **$1.411 \times 10^{-7}$** | **$2.754 \times 10^{-11}$** | **$6.566 \times 10^{-10}$** | **$3.036 \times 10^{-10}$** |
| **years of education** | ***β*** | -0.05227565 | -0.03223422 | -0.07457150 | -0.08864574 | -0.05570660 |
| | **Std. Error** | 0.05605588 | 0.05396724 | 0.05228354 | 0.05722661 | 0.06033889 |
| | **95% CI** | -0.163 – 0.058 | -0.138 – 0.074 | -0.178 – 0.028 | -0.201 – 0.024 | -0.175 – 0.063 |
| | ***t*** | -0.9325632 | -0.5972924 | -1.4262900 | -1.5490301 | -0.9232288 |
| | ***p*** | 0.3518881 | 0.5508184 | 0.1549572 | 0.1225545 | 0.3567216 |
| **gender [male vs. female]** | ***β*** | -0.9238238 | -1.0935897 | -0.9793131 | -1.1213333 | -0.9137877 |
| | **Std. Error** | 0.2923866 | 0.2719083 | 0.2669394 | 0.2752757 | 0.2835604 |
| | **95% CI** | -1.500 – -0.348 | -1.629 – -0.558 | -1.505 – -0.454 | -1.663 – -0.579 | -1.472 – -0.355 |
| | ***t*** | -3.159597 | -4.021907 | -3.668673 | -4.073492 | -3.222550 |
| | ***p*** | **$1.762 \times 10^{-3}$** | **$7.519 \times 10^{-5}$** | **$2.945 \times 10^{-4}$** | **$6.104 \times 10^{-5}$** | **$1.428 \times 10^{-3}$** |
| **RMSE** | | 1.746853 | 2.426911 | 2.461695 | 2.264960 | 1.985310 |
| **df** | | 267 | 267 | 266 | 268 | 267 |

*Note*. Runs correspond to test folds used, i.e. run 1 corresponds to a LM with fold 1 as the test dataset and the remaining data (folds 2, 3, 4 and 5) as training data. Bold *p*-values are significant at an alpha level of .05. All *p*-values are uncorrected. RMSE = Root Mean Squared Error, CI = Confidence Interval.

### Table S7. Results of the LMM for reading times.

| Predictor | Estimate | Std. Error | CI | t | p | | df |
|---|---|---|---|---|---|---|---|
| reading time of previous trial (log-transf.) | 0.276268 | 0.001784 | 0.272771 – 0.279764 | 154.849742 | $< 1.307 \times 10^{-97}$ | * | 158637.71 |
| mean d-prime singletasks | 0.033253 | 0.017540 | -0.001300 – 0.067806 | 1.895787 | $6.659 \times 10^{-2}$ | | 239.31 |
| mean comprehension question performance | 0.003828 | 0.001243 | 0.001380 – 0.006276 | 3.080216 | $2.970 \times 10^{-3}$ | * | 239.47 |
| de-meaned comprehension question performance | -0.000479 | 0.000051 | -0.000578 – -0.000379 | -9.410071 | $1.291 \times 10^{-20}$ | * | 157527.98 |
| word frequency | 2.087721 | 0.451323 | 1.201794 – 2.973648 | 4.625781 | $6.534 \times 10^{-6}$ | * | 794.04 |
| word length | 0.011112 | 0.000489 | 0.010152 – 0.012072 | 22.703634 | $1.307 \times 10^{-97}$ | * | 1481.42 |
| word entropy | 0.001593 | 0.000886 | -0.000145 – 0.003330 | 1.796885 | $7.664 \times 10^{-2}$ | | 9812.21 |
| block number | -0.008956 | 0.000118 | -0.009187 – -0.008724 | -75.763375 | $< 1.307 \times 10^{-97}$ | * | 156255.40 |
| trial number | -0.000385 | 0.000008 | -0.000401 – -0.000369 | -47.770204 | $< 1.307 \times 10^{-97}$ | * | 23165.89 |
| recording location [online vs. lab] | -0.278456 | 0.028593 | -0.334782 – -0.222129 | -9.738450 | $9.606 \times 10^{-19}$ | * | 239.62 |
| surprisal | 0.001293 | 0.000172 | 0.000955 – 0.001631 | 7.503810 | $1.581 \times 10^{-13}$ | * | 3333.63 |
| age | 0.006313 | 0.000807 | 0.004724 – 0.007903 | 7.825449 | $2.919 \times 10^{-13}$ | * | 239.40 |
| ASC score | 0.012846 | 0.005625 | 0.001765 – 0.023926 | 2.283781 | $2.791 \times 10^{-2}$ | * | 239.32 |
| surprisal × age | 0.000061 | 0.000004 | 0.000052 – 0.000069 | 13.886055 | $2.447 \times 10^{-43}$ | * | 161362.62 |
| surprisal × ASC score | -0.000193 | 0.000032 | -0.000256 – -0.000130 | -5.999267 | $3.250 \times 10^{-9}$ | * | 161141.90 |
| age × ASC score | -0.000148 | 0.000347 | -0.000831 – 0.000536 | -0.425008 | $6.712 \times 10^{-1}$ | | 239.28 |
| surprisal × age × ASC score | -0.000008 | 0.000002 | -0.000011 – -0.000004 | -4.004004 | $8.627 \times 10^{-5}$ | * | 160734.68 |
| N | 247 | | | | | | |
| Intra-class Correlation (ICC) | 0.48 | | | | | | |
| Marginal $R^2$ / Conditional $R^2$ | 0.489 / 0.736 | | | | | | |

*Note.* All continuous predictors were mean-centred. Degrees of freedom for *p*-values, standard errors and confidence intervals (CI) were computed using Satterthwaite's approximation. All *p*-values reported here are FDR-corrected and were computed using ANOVAs with type III sum of squares. Results that are significant on an alpha-level of 0.05 are marked with a star.

***Control analysis: 5-fold cross-validation of the age × surprisal × ASC interaction***

To probe the robustness and generalisability of the interaction of age, surprisal and ASC scores, we again employed 5-fold cross-validation (21). Across the five runs, the direction of the interaction remained stable, with effects from three of the five models reaching significance (see Fig. S3 and Table S8. The mean RMSE was 0.307 ± 0.030, which indicates an average deviation between predicted and test data of approximately 0.31 log-units in reading time, which corresponds to an average multiplicative prediction error of roughly 35.9%. Across folds, our models showed strong and stable explanatory power, with a mean conditional $R^2$ of 0.737 ± 0.011 and a mean marginal $R^2$ of 0.489 ± 0.023. This indicates that the full mixed model explained approximately 74% of the variance in reading times overall, while the fixed effects alone accounted for roughly 49% of the variance.

**Table S8.** Results of the 5-fold cross-validation for the 3-way interaction effect of surprisal, age and ASC score on reading time (log-transformed).

| | Estimate | Std. Error | 95% CI | t | p | df | RMSE |
|---|---|---|---|---|---|---|---|
| **run 1** ($N_{test}$ = 50, $N_{training}$ = 197) | -0.000004 | 0.000002 | -0.000008 – 0.0000007 | -1.656096 | $9.770\times10^{-2}$ | 126240.1 | 0.286062 |
| **run 2** ($N_{test}$ = 50, $N_{training}$ = 197) | -0.000010 | 0.000002 | -0.000013 – -0.0000053 | -4.411248 | **$1.029\times10^{-5}$** | 128200.2 | 0.355946 |
| **run 3** ($N_{test}$ = 49, $N_{training}$ = 198) | -0.000010 | 0.000002 | -0.0000144 – -0.0000064 | -5.050222 | **$4.419\text{x}10^{-7}$** | 130334.9 | 0.306848 |
| **run 4** ($N_{test}$ = 49, $N_{training}$ = 198) | -0.000003 | 0.000002 | -0.0000076 – 0.0000009 | -1.542660 | $1.229\times10^{-1}$ | 128856.7 | 0.278316 |
| **run 5** ($N_{test}$ = 49, $N_{training}$ = 198) | -0.000011 | 0.000002 | -0.0000146 – -0.0000064 | -5.051006 | **$4.401\times10^{-7}$** | 129100.1 | 0.308288 |

*Note*. Runs correspond to test folds used, i.e. run 1 corresponds to a LMM with fold 1 as the test dataset and the remaining data (folds 2, 3, 4 and 5) as training data. All continuous predictors were mean-centred. Degrees of freedom for *p*-values, standard errors and confidence intervals (CI) were computed using Satterthwaite's approximation. All *p*-values reported here are uncorrected and were computed using ANOVAs with type III sum of squares. Bold *p*-values are significant at an alpha level of .05. RMSE = Root Mean Squared Error, CI = Confidence Interval.

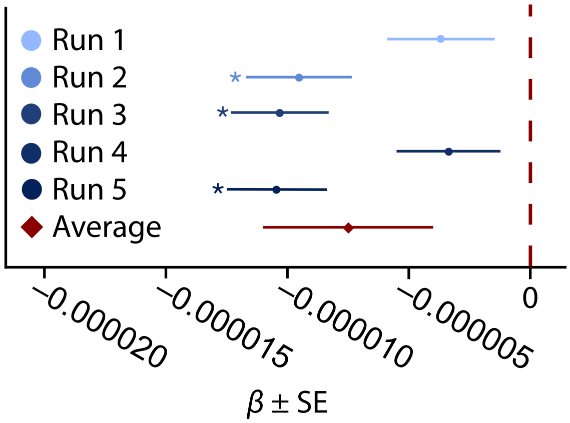


**Figure S3.** Results of the 5-fold cross-validation for the 3-way interaction effect of surprisal, age and ASC score on reading time (log-transformed). Presented are estimates ± standard errors (SE) per run, with each run comprising 80% of the full dataset (N = 247) as training data. Average estimates were calculated across runs. To pool SEs, we averaged the squared SEs (variances) from each fold, added the variance of the estimates across folds adjusted for sample size, and then took the square root to reflect total uncertainty. Effects that are significant on an alpha level of .05 are marked with a star (p-values uncorrected).

#### ***Control analysis: Contribution of AQ and SPQ subscales to the age × surprisal × ASC interaction***

In a control analysis, we examined which AQ and SPQ subscales most strongly contributed to the significant 3-way interaction of age, surprisal and ASC score. Specifically, we asked whether the observed effect is primarily driven by cognitive-perceptual traits or whether age-related variation in communication and social functioning accounts for the observed pattern.

To address this, we fit the linear mixed-effects model reported in the main text (see analysis of reading times), replacing the composite ASC score with individual subscale scores. We selected the subscales with the highest loadings on the composite score (AQ: Social Skill [SS], Communication [CM], Imagination [I], and Repetitive and Restrictive Behaviour [RBB]; SPQ: Ideas of Reference [IR] and Unusual Perceptual Experiences [UPE]). This resulted in six separate models, each including one subscale as the trait predictor of interest.

The critical three-way interaction (Age × Surprisal × Subscale) remained significant in five of the six models, failing to reach significance only for the AQ subscale Repetitive and Restrictive Behaviour (RBB). Across models, two robust patterns emerged (Fig. S4). First, considering the sign inversion required for SPQ subscales due to their negative loadings on the composite ASC score, the direction of effects was highly consistent: older adults showed stronger surprisal effects, particularly when exhibiting lower subscale scores indicative of better social-communicative functioning and reduced tendencies toward schizotypal perception and cognition. Second, and more importantly, modulation of surprisal was not restricted to subscales indexing communication and social behaviour. Comparable effects were also observed for subscales reflecting cognitive-perceptual variation. This indicates that the observed interaction cannot be explained solely by differences in communicative style, but instead reflects broader cognitive-perceptual trait differences, which further validates the interpretation of the ASC as indexing cognitive-perceptual style.

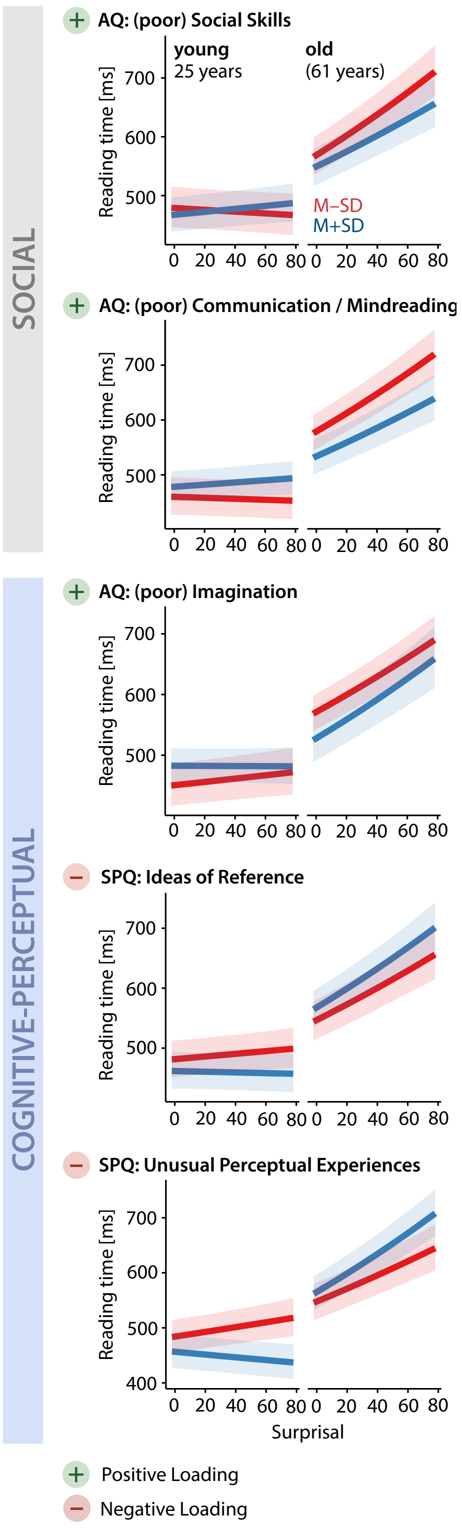


**Figure S4.** Estimated marginal means for 3-way interactions of AQ and SPQ subscales with age and surprisal on self-paced reading times (N = 247). Only results from models showing a significant three-way interaction are displayed (α = 0.05, FDR-corrected).
